# Multi-omics characterization of lymphedema-induced adipose tissue from breast cancer patients

**DOI:** 10.1101/2023.11.03.565593

**Authors:** Sinem Karaman, Satu Lehti, Cheng Zhang, Marja-Riitta Taskinen, Reijo Käkelä, Adil Mardinoglu, Håkan Brorson, Kari Alitalo, Riikka Kivelä

## Abstract

**Objective:** Secondary lymphedema (LE) following breast cancer surgery is a life-long complication, which currently has no cure. LE induces significant regional adipose tissue deposition, requiring liposuction as a treatment. Here, we aimed to elucidate the transcriptional, metabolomic, and lipidomic signature of the adipose tissue developed due to the surgery-induced LE in short- and long-term LE patients, and compared the transcriptomic landscape in LE to the obesity-induced adipose tissue.

**Methods:** Adipose tissue biopsies were obtained from breast cancer-operated females with LE from the affected and non-affected arms (n=20 patients). To decipher molecular properties of the LE adipose tissue, we performed RNA sequencing, metabolomics, and lipidomics combined with bioinformatics analyses.

**Results:** Integrative analysis of functional genomics revealed that inflammatory response, cell chemotaxis and angiogenesis were upregulated biological processes in the LE arm, indicating a sustained inflammation in the edematous adipose tissue, whereas, epidermal differentiation, cell-cell junction organization, water homeostasis and neurogenesis were, in turn, downregulated in the LE arm. Surprisingly, only few genes were found to be the same in the LE-induced and the obesity-induced adipose tissue expansion, indicating a different type of adipose tissue development in these two diseases. In metabolomics analysis, the concentration of a branched-chain amino acid valine was found to be reduced in the edematous arm together with downregulation of mRNA levels of its transporter *SLC6A15*. Lipidomics analyses did not show any significant differences between the diseased and healthy arm, suggesting that diet affects the lipid composition of the adipose tissue more than the LE.

**Conclusions:** Our results provide a detailed molecular characterization of adipose tissue in secondary LE of breast cancer patients vs individuals with obesity. The results show distinct differences in transcriptomic signatures between LE patients vs individuals with obesity, but only minor differences in metabolome and lipidome between the diseased and the healthy arm.

## INTRODUCTION

Lymphedema (LE) is an excessive accumulation of a protein-rich extracellular fluid (lymph) in the interstitial compartment due to defective lymphatic function (reviewed in ^1,2^). The accumulation of fluid elicits an inflammatory reaction, induces fibrosis and adipose tissue accumulation, and leads to impaired wound healing and immune responses ^3,4^. The prevalence estimates of chronic LE are between 1.33 to 1.44 per 1000 individuals. However, unreported cases are common, thus it is thought that the reported prevalence is an underestimate ^5^. LE is categorized into two subclasses according to the underlying pathophysiology: primary LE, also known as hereditary LE, and secondary (acquired) LE ^1^. While primary LE is congenital, secondary LE is caused by filariasis or obliteration of the lymphatic vasculature or lymph nodes by trauma, surgery, radiation therapy, or infection ^3,6^.

Secondary LE is more common, and its pathophysiology has been more comprehensively studied (reviewed in ^7-9^). Filariasis is the most common cause of secondary LE, which is caused by the *Wuchereria bancrofti* or *Brugia malayi/timori* parasite infection, and it is observed mainly in tropical regions, affecting more than 100 million people worldwide ^6^. In contrast, breast cancer-related LE is the most common form of LE in the developed countries ^10^, predominantly related to cancer treatment: the surgical removal of lymph nodes and radiotherapy. In the aftermath of the surgical procedure to remove the lymph nodes, which is often followed by irradiation to kill remaining tumor cells, the lymphatic vasculature is commonly damaged beyond repair. This initially leads to a dilation of the lymphatic vasculature, followed by increased lymphatic endothelial cell proliferation but ultimately leads to lymphatic valve malfunction, fibrosis of smooth muscle cells in the walls of the lymphatics, and LE. To date, there is no “silver bullet” for a treatment or cure for LE. The available treatment approaches aim at reducing the symptoms and mostly rely on physiotherapy, complete decongestive therapy (CDT) comprising of manual drainage, bandaging, exercises, flat-knitted compression garments and skin care ^11^ or controlled compression therapy by restrictive bandaging ^12^. Surgical alternatives include liposuction of the edematous limb ^12-16^ and microvascular reconstruction, involving lymphatico-venous or lymphatico-venous-lymphatic shunts ^17-22^, lymph node transfer ^23-25^, or lymph vessel transplantation ^26-29^.

LE patients often present with local adipose tissue overgrowth (reviewed in ^1,2^). Arm LE is a common complication after surgical removal of axillary lymph nodes for breast cancer treatment ^30-33^. Neither CDT nor microsurgical reconstruction can be used in later stages of LE as none of the techniques can remove the hypertrophied adipose tissue that occurs in response to lymph stasis and inflammation ^30,31,33^. In later stages of non-pitting LE, which does not respond to conservative treatment, liposuction, combined with postoperative CCT, gives a complete reduction of the excess volume ^12-16^.

Increased adiposity in mouse models of obesity has been shown to result in lymphatic dysfunction ^34,35^. On the other hand, LE and leaky lymphatic vessels have been shown to induce adipogenesis and adipose tissue accumulation also in mouse models ^36,37^. Adipogenesis in response to lymphatic fluid stasis is associated with a marked mononuclear cell inflammatory response and up-regulation of the expression of adipocyte differentiation markers ^38^. It seems that the underlying pathophysiology of LE drives adipose-derived stem cells toward adipogenic differentiation ^39^. These data support the clinical finding that leaky lymphatics and lymph stasis in LE patients can result in adipose tissue accumulation in the affected region.

The factors involved in LE-induced adipose tissue accumulation are poorly known. Furthermore, it is not known if adipose tissue growth in LE and obesity follow similar mechanisms. So far, the treatments are only aimed at alleviating LE symptoms, without targeting the underlying causes. Here, we aimed to obtain an unbiased insight into the transcriptional, metabolomic, and lipidomic changes in LE-induced adipose tissue in human patients. We found that the transcriptomic landscape of the LE-induced adipose tissue is highly different from that of the unaffected arm, and it markedly differed from obesity-induced adipose tissue. In contrast, the metabolomic and lipidomic profile of the adipose tissue in the LE arm did not differ much from the healthy arm, suggesting that that the individual diet and systemic pool of metabolites are stronger determinants of the metabolome and lipidome of the local adipose tissue than the disease.

## MATERIALS AND METHODS

### Patients and sample collection, consent, and ethical permit

Subcutaneous adipose tissue was obtained from the arms of 20 female patients undergoing liposuction for chronic LE induced by breast cancer treatment. From these, samples of 16 patients were available for lipidomic analysis. Control subcutaneous adipose tissue was collected from the unaffected arm of the same individuals. Duration of LE was determined as the time from surgery to liposuction, and the patients were divided into short- (≤3 years) and long- (≥14 years) term LE groups (n=10/7 in short-term group and 10/9 in the long-term group were available for lipidomics). Work described has been carried out in accordance with The Code of Ethics of the World Medical Association (Declaration of Helsinki) and informed consent was obtained for tissue collection. All patients were given general anesthetics, and before the start of liposuction, a biopsy was taken after a 1.5-cm incision on the medial aspect of the elbow. The ethical permit for the collection of patient samples is held by Dr. Håkan Brorson and was approved by the Regional Ethical Review Board in Lund, Lund, Sweden (503/2006, 45/2011).

### RNA sequencing and analysis

Total RNA was extracted using Trizol and purified with RNeasy Mini kit (Qiagen). Total RNA was subjected to quality control with Agilent Bioanalyzer according to the manufacturer’s instructions. To construct libraries suitable for Illumina sequencing the Illumina TruSeq Stranded mRNA Sample preparation protocol which includes cDNA synthesis, ligation of adapters and amplification of indexed libraries was used. The yield and quality of the amplified libraries were analyzed using Qubit by Thermo Fisher and the Agilent Tapestation. The indexed cDNA libraries were normalized and combined, and the pools were sequenced on the Illumina HiSeq 2000 for a 50-cycle v3 sequencing run generating 50 bp single-end reads. Basecalling and demultiplexing were performed using CASAVA software with default settings generating Fastq files for further downstream mapping and analysis. Raw counts were aligned using STAR and counting the reads in gene bodies were defined using a UCSC gtf file. The counts were normalized to counts per million and after TMM normalization in the EdgeR package. Differential expression was analyzed using the DEseq2 package in R.

Venn diagrams comparing the significantly changed transcripts in short and long term LE patients were made using the online Venn diagram tool at http://bioinformatics.psb.ugent.be/webtools/Venn/. ShinyGo was used to analyze gene ontologies and functional genomics from the RNAseq data (http://bioinformatics.sdstate.edu/go/).

### Quantitative real-time PCR

RNA was extracted from frozen adipose tissue pieces (50-100 mg) with the RNeasy Lipid tissue Mini Kit (Qiagen) following the manufacturer’s instructions. cDNA was transcribed from 500 ng RNA template, using the High-Capacity Reverse Transcription kit (Applied Biosystems). The RT-qPCR was performed using a BIO-RAD C1000 Thermal cycler according to a standardized protocol (BIO-RAD), with Roche SYBR green. *RPLP0* (*36B4*) was used as internal control and fold changes of gene expression were calculated using the DeltaDeltaCt method. The primer sequences used can be found in **Supplemental Table 1**.

### Reporter metabolite analysis

Reporter metabolite analysis was run on the data from RNA seq using previously published pipeline ^40^. In brief, this method retrieves a gene-metabolite network from the genome-scale metabolic models (GEMs) and identifies key metabolite hubs with the outstanding amount of DE genes that are obtained as described above directly connecting to them. A previously published high-quality GEMs for adipose tissue ^41^ were used as input for the reporter metabolite analysis.

### Comparison to obesity dataset

The differentially expressed genes (DEGs) between white adipose tissues obtained from lean subjects and subjects with obesity (all females) were retrieved from a previous study ^41^, and the enriched gene ontology (GO) term network of the overlapped genes was visualized using cytoscape with the external package EnrichmentMap ^42^.

### Metabolomics

Metabolomics analyses were carried out at the Biocenter Finland and HiLIFE supported Metabolomics Unit, Institute for Molecular Medicine Finland FIMM, HiLIFE, University of Helsinki. Approximately 20 mg of frozen adipose tissue per sample was analyzed and normalized to the exact weight according to the method described ^43^.

### Lipidomics

Seven pairs of samples from short-term LE and nine pairs of samples from long-term LE were available for lipidomic analysis. Lipidomic analyses of adipose tissue pieces (50-100 mg) included shotgun electrospray ionization tandem mass spectrometry (ESI-MS/MS) of triacylglycerol (TAG), phosphatidylcholine (PC) and sphingomyelin (SM) species, and gas chromatography of fatty acids (FA) (FID and MS detections for quantification and structure confirmation, respectively). These analyses were carried out as described ^44^, except, in the present work, the equipment recording electron impact mass spectra for FA structure confirmation was Shimadzu GCMS-QP2010 Ultra (Shimadzu Scientific Instruments, Kyoto, Japan).

### Analysis of lipidomics data

Instead of separately presenting fold change values for the relative concentrations of numerous individual FA and lipid species in adipose tissues from the LE and healthy arms, the FA compositions in total lipids and lipid species compositions in each lipid class were compared by Principal Component Analysis (PCA). The magnitude of the compositional difference on PC1 and PC2 scores was normalized to the average compositional difference of the two arms in the 16 individuals and illustrated in heatmap format.

## RESULTS

Paired subcutaneous adipose tissue samples were collected from both forearms of 20 breast cancer patients who had developed unilateral post-surgical LE. The patients were divided into short and long-term LE groups (short ≤3 years and long ≥14 years, respectively). The collected adipose tissue samples were then used for transcriptomic, metabolomic, and lipidomic analyses. The study design, patient demographic data, and clinical characteristics of the patients who underwent breast cancer-related surgery are shown in **Figure 1**.

**Figure 1.**
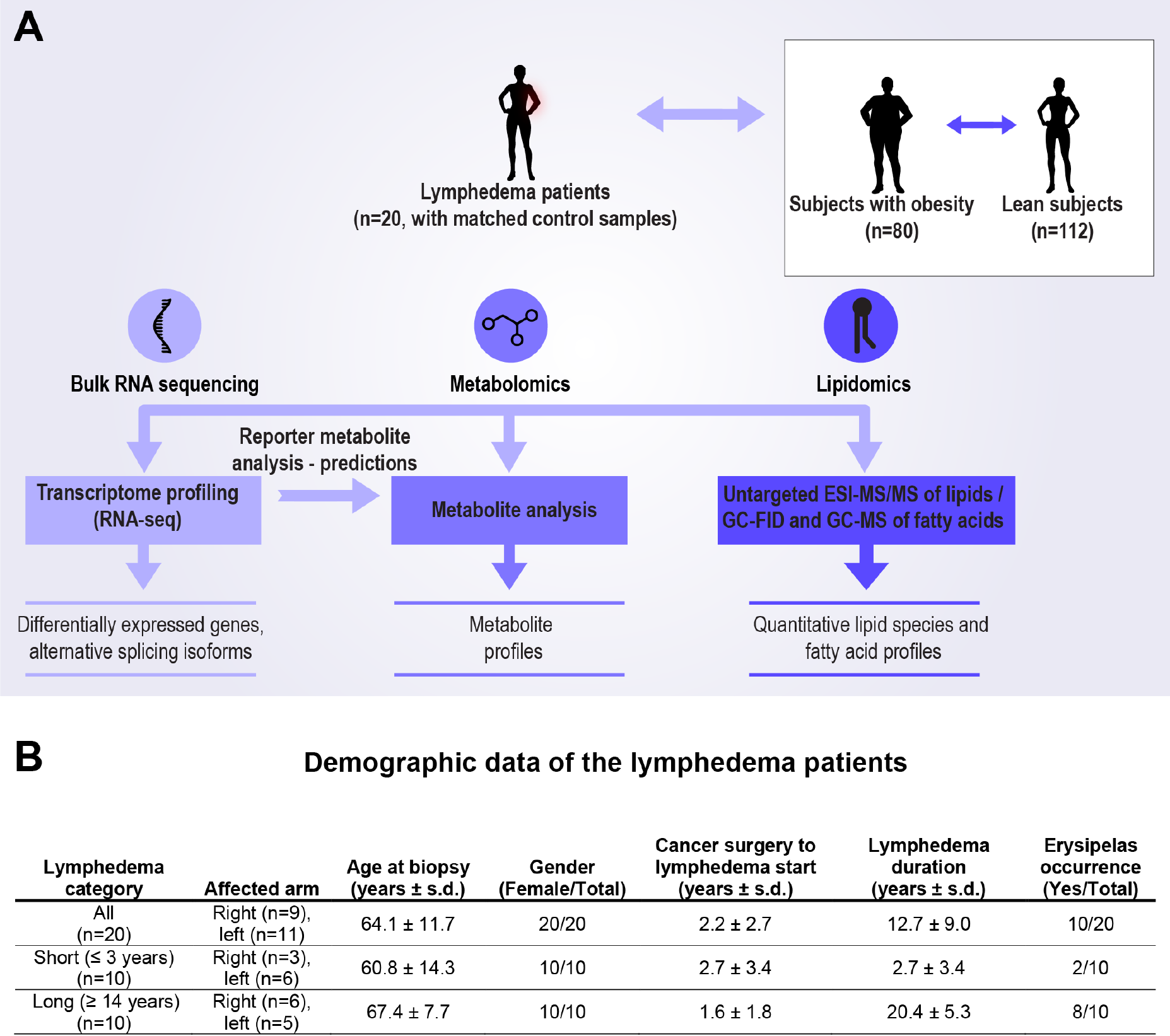
Study design and demographic data of the patients. (**A**) The multi-omics approach to analyze adipose tissue biopsies obtained from female breast cancer patients who have developed lymphedema (LE) after surgery. Transcriptomics data from the adipose tissue of the edematous arm has been compared to the differentially expressed genes of adipose tissues of subjects with obesity. (**B**) Demographic data of the patients.

### Altered adipose tissue gene signature in LE

To obtain an unbiased insight into the transcriptional changes associated with LE in adipose tissue of the lymphedematous arm, we performed global gene expression analyses with RNA sequencing using the unaffected arm as a control. We identified a total of 1473 transcripts that were dysregulated in all LE patients, of which 830 were upregulated and 643 downregulated (**Supplemental Data Table 1**). The most upregulated genes in the LE arm included genes related to cholesterol metabolism (e.g. *CETP, ABCG1*) and to inflammation (e.g. *VEGFC, VCAM1*; **Figure 2A**). Cholesteryl ester transfer protein (*CETP*), a plasma protein involved in cholesteryl ester and TAG transfer between LDL and HDL, was the most significantly upregulated gene in LE. The most significantly repressed gene was dermcidin (*DCD*), an important antimicrobial gene produced by the sweat glands of the skin ^45^. Among the most downregulated genes was also *SLC6A15*, a branched-chain amino acid transporter. In addition, several keratins were downregulated in the adipose tissue samples from the LE arms.

**Figure 2.**
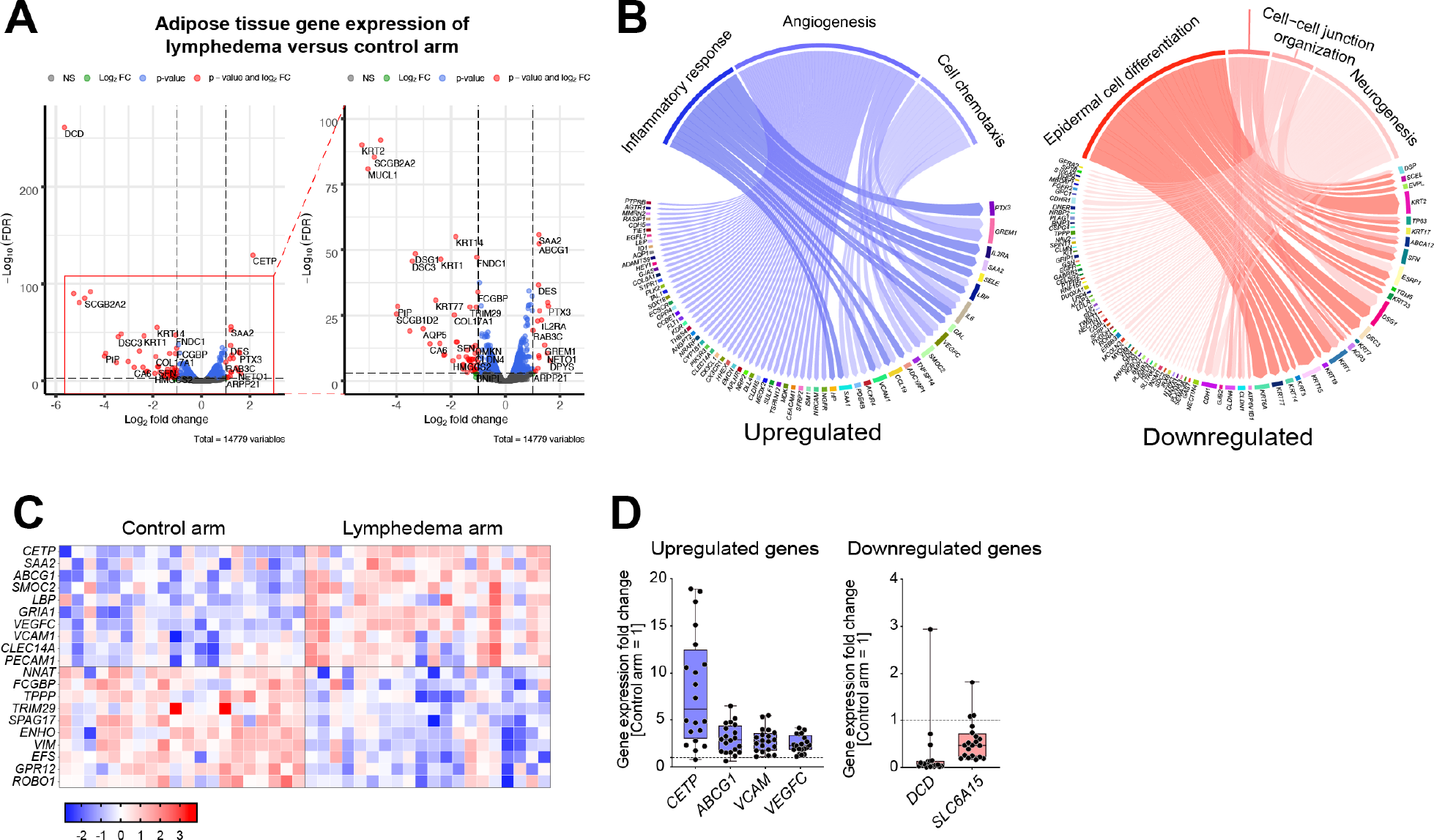
RNA sequencing of adipose tissue from the lymphedema and the control arms. (**A**) Volcano plots of all differentially expressed genes between the LE arm and the control arm (FDR cut-off = 0.001, log_10_ FC <-0.25 or >0.25, n=20), and a zoom in for the transcripts shown in the red rectangle. (**B**) Significantly up- and downregulated GO terms and selected genes within the GO terms suggest an upregulation in inflammatory response, angiogenesis and cell chemotaxis pathways, whereas epidermal cell differentiation, water homeostasis, cell-cell junction organization and neurogenesis pathways were significantly downregulated. (**C**) Heatmap of selected genes up- or downregulated in the lymphedema arm compared to the control arm. (**D**) qPCR validation of selected genes. Dashed line indicates the expression level in the control arm (=1).

GO enrichment analysis of the upregulated genes showed that inflammatory response, cell chemotaxis and angiogenesis were among the most affected biological processes in the adipose tissue from the LE arm, indicating a sustained inflammation that may increase blood and lymph flow in the edematous adipose tissue (**Figure 2B**). Genes related to epidermal differentiation, cell-cell junction organization, water homeostasis, and neurogenesis were found to be repressed in the LE vs. control arm (**Figure 2B**). Heatmap of selected genes’ Z-scores and the validation of gene expression changes with qPCR are shown in **Figures 2C and 2D**, respectively.

### Different transcriptomic landscapes in short-term and long-term LE

Next, we studied if the LE adipose tissue transcriptome is affected by the duration of the disease. The patients were divided into short and long-term LE groups. Comparison of the differentially expressed genes between these subgroups demonstrated an overlap in both up- and downregulated genes between the groups, but also a significant number of genes unique for each group. This indicates that the transcriptomic landscape in LE-derived adipose tissue continues to evolve after the initial development of the disease. The Venn diagrams comparing the groups are shown in **Figures 3A and 3B**. Gene ontology analysis of the genes unique for each subgroup revealed that in the short-term LE group, angiogenesis, vasculature development, and immune response were the most upregulated pathways (**Figure 3C**), indicating an immune response and an attempt to restore blood and lymphatic circulation. In the long-term LE group, muscle contraction and extracellular matrix organization were the most activated pathways, followed by vasculature development (**Figure 3D**). The most repressed pathways in the short-term LE were the establishment of skin barrier, water homeostasis, keratinization, and neurogenesis (**Figure 3E**), whereas, in the long-term LE tissues, lipid metabolism, fat cell differentiation and response to insulin were highly downregulated (**Figure 3F**). These results indicate that LE first affects skin development and function and, at later stages, mainly adipose tissue metabolism.

**Figure 3.**
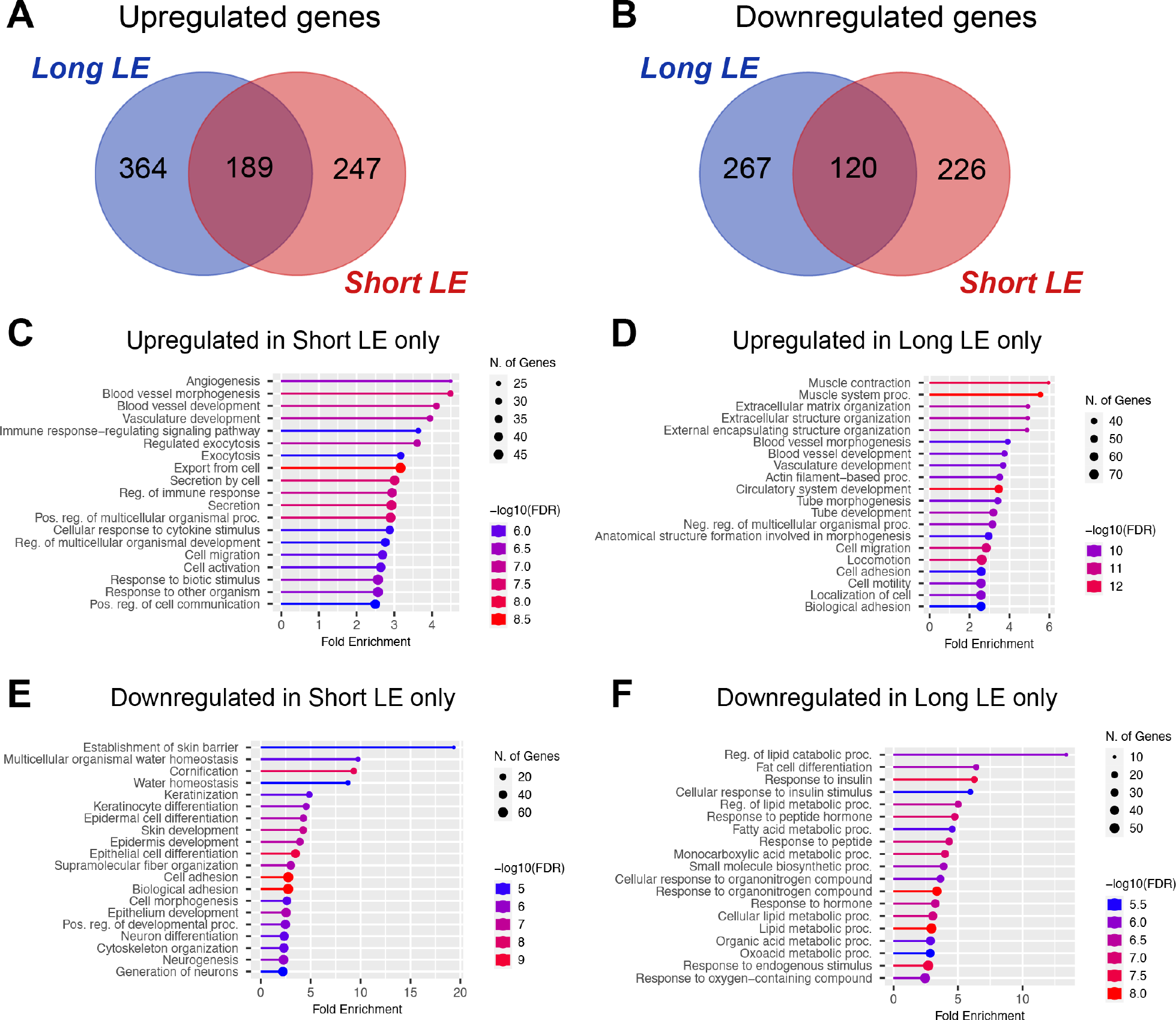
Comparison of the short-term and long-term LE. (**A, B**) Venn-diagrams showing genes significantly up (**A**) or downregulated (**B**) in short-term (≤3 years) and long-term (≥14 years) LE (FDR cut-off = 0.01, log_10_ FC <-0.25 or >0.25, n=10 versus 10). (**C**) GO term analysis of the 247 genes upregulated only during short-term LE. (**D**) GO term analysis of the 364 genes upregulated only during long-term LE. (**E**) GO term analysis of the 226 genes downregulated only during short-term LE. (**F**) GO term analysis of the 267 genes downregulated only during long-term LE.

### Comparison of adipose tissues in LE and obesity

To compare the properties of adipose tissue from LE patients to those of patients with obesity (BMI≥30), we used data from subcutaneous adipose tissues of females with and without obesity (n=80 with obesity and n=112 lean controls) from a previously published cohort ^41^.

Surprisingly, we found very little overlap between adipose tissue transcriptomic changes in obesity versus LE, as out of the 1682 differentially expressed genes (DEG) between subjects with obesity and lean subjects and 1473 DEGs between the LE and healthy arm, only 225 genes were shared between the two conditions (**Figure 4A**). This suggested major differences in the development and characteristics of these two adipose tissue types. The GO enrichment analysis showed that the genes that were common to both conditions were related to extracellular matrix organization, cell proliferation and migration, and circulatory system development (**Figure 4B and 4C**).

**Figure 4.**
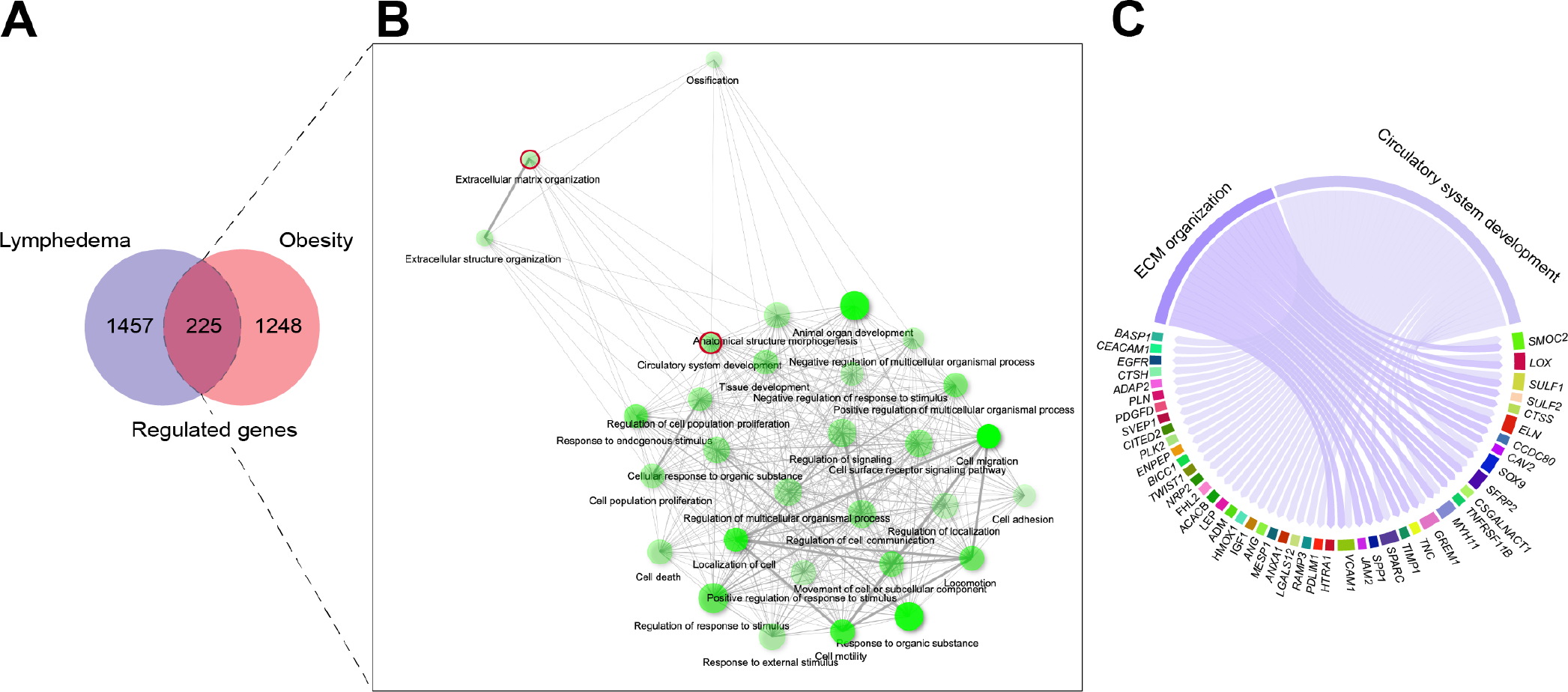
Differential transcriptomic changes during the subcutaneous adipose tissue enlargement in lymphedema versus obesity. (**A**) A Venn-diagram showing the number of affected genes in lymphedema adipose tissue compared to adipose tissue from subjects with obesity. Only 225 genes were significantly co-regulated in these two conditions. (**B**) GO term networks of the 225 genes. (**C**) Genes that fall under the GO term extracellular matrix (ECM) organization and circulatory system development.

### Reporter metabolite predictions and metabolomics results

In order to have a better understanding of potential changes in metabolites, we performed a reporter metabolite analysis by using the RNA sequencing data as input, as previously described ^40^. Based on the changes in gene expression, this analysis predicted an upregulation of nucleotide metabolism and a downregulation of amino acid metabolism including branched-chain amino acids (leucine, isoleucine, and valine) in LE adipose tissue vs. the healthy arm (**Figure 5A**). To compare the reporter metabolite predictions with the actual metabolite content of the tissue, we performed metabolomics from the frozen adipose tissue samples (**Figure 5B**). Overall, we did not observe major differences in the concentrations of the analyzed metabolites between the control and the lymphedematous arms. The only significant difference was in valine concentration, which was also predicted to be reduced by the reporter metabolite analyses. We then returned to the RNA sequencing data and checked the expression levels of branched-chain amino acid transporters. Out of the three transporters *SLC6A15, SLC6A14*, and *SLC6A19* that are important for valine metabolism, we observed that *SLC6A15* was one of the most downregulated genes in our RNAseq dataset. These results were further validated by qPCR using the subcutaneous adipose tissue samples (**Figure 2D**).

**Figure 5.**
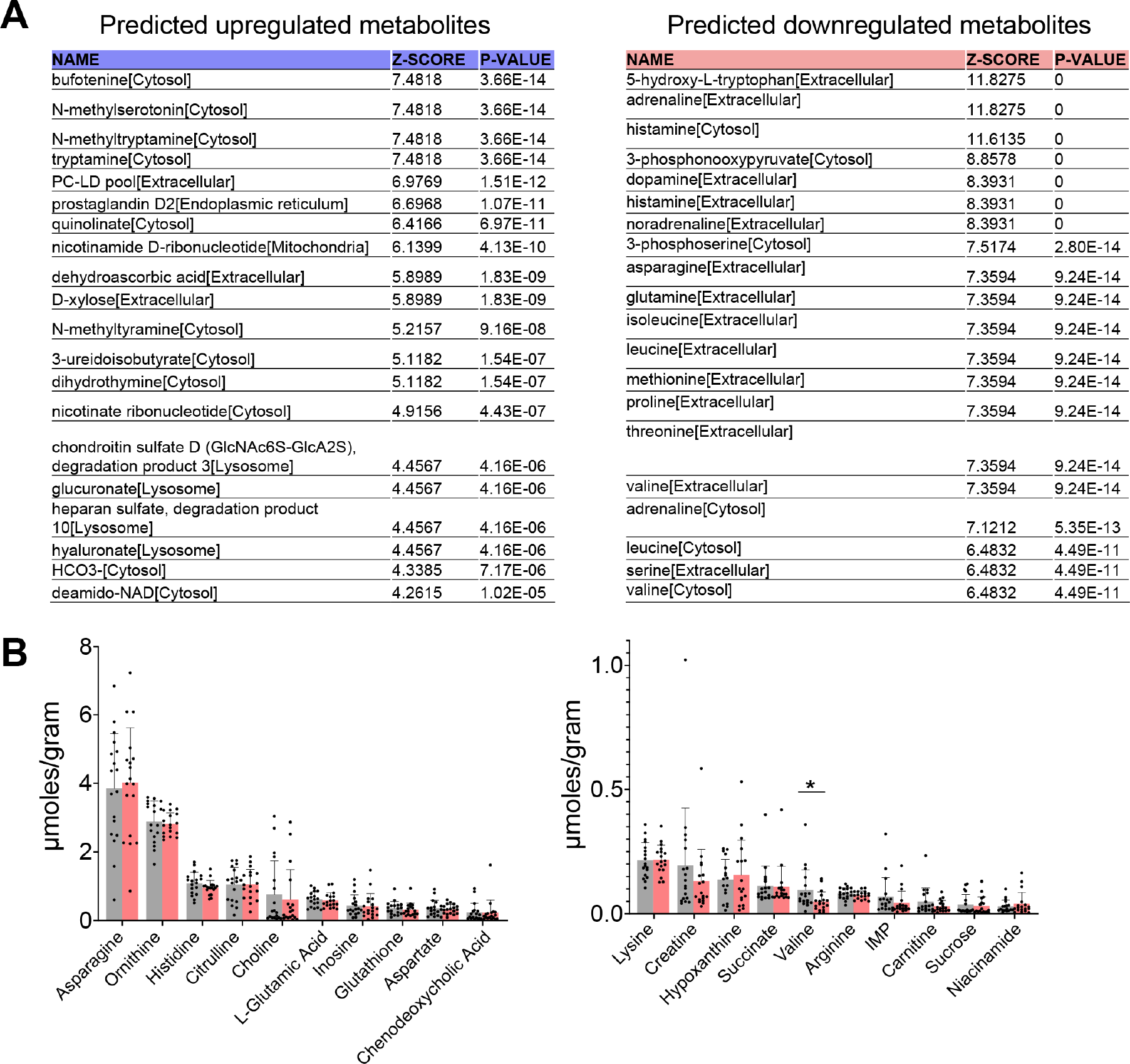
Predicted reporter metabolites and metabolomics analysis. (**A**) Top 20 predicted upregulated and downregulated reporter metabolites based on the RNA sequencing data suggest upregulation of nucleoside metabolism and downregulation of amino acid metabolism in the LE adipose tissue. (**B**) Targeted analysis of metabolites between the adipose tissues of control and lymphedema arm of the same patient shows mostly unchanged metabolites with the exception of lower valine levels (* FDR<0.05).

### Lipidomics results

To analyze the effect of LE on adipose tissue lipid species composition, we next performed untargeted lipidomics with mass spectrometry and FA profiling by gas chromatography. We were able to detect the main species of TAG, PC, and SM, and the FA profile of total lipids (**Figure 6**). The main finding was that the lipid species and FA profiles showed large variation between different patients but not consistently between LE versus control arms, suggesting that the diet has a bigger role than the disease in modulating the lipid and FA composition of the adipose tissue (**Figure 6A-E**). When a relatively large compositional difference between the two arms was found, the patient had either short- or long-term LE (**Figure 6A**). In several of these individuals, a large difference between the arms in the composition of storage lipid (TAG) was accompanied by a large difference also in membrane lipids (PC and SM). Since the individual FA or lipid species responsible for the compositional difference varied between the patients (the LE samples had shifted from their control pairs to different directions on the PCA plots), the potential metabolic aberrations behind were individual.

**Figure 6.**
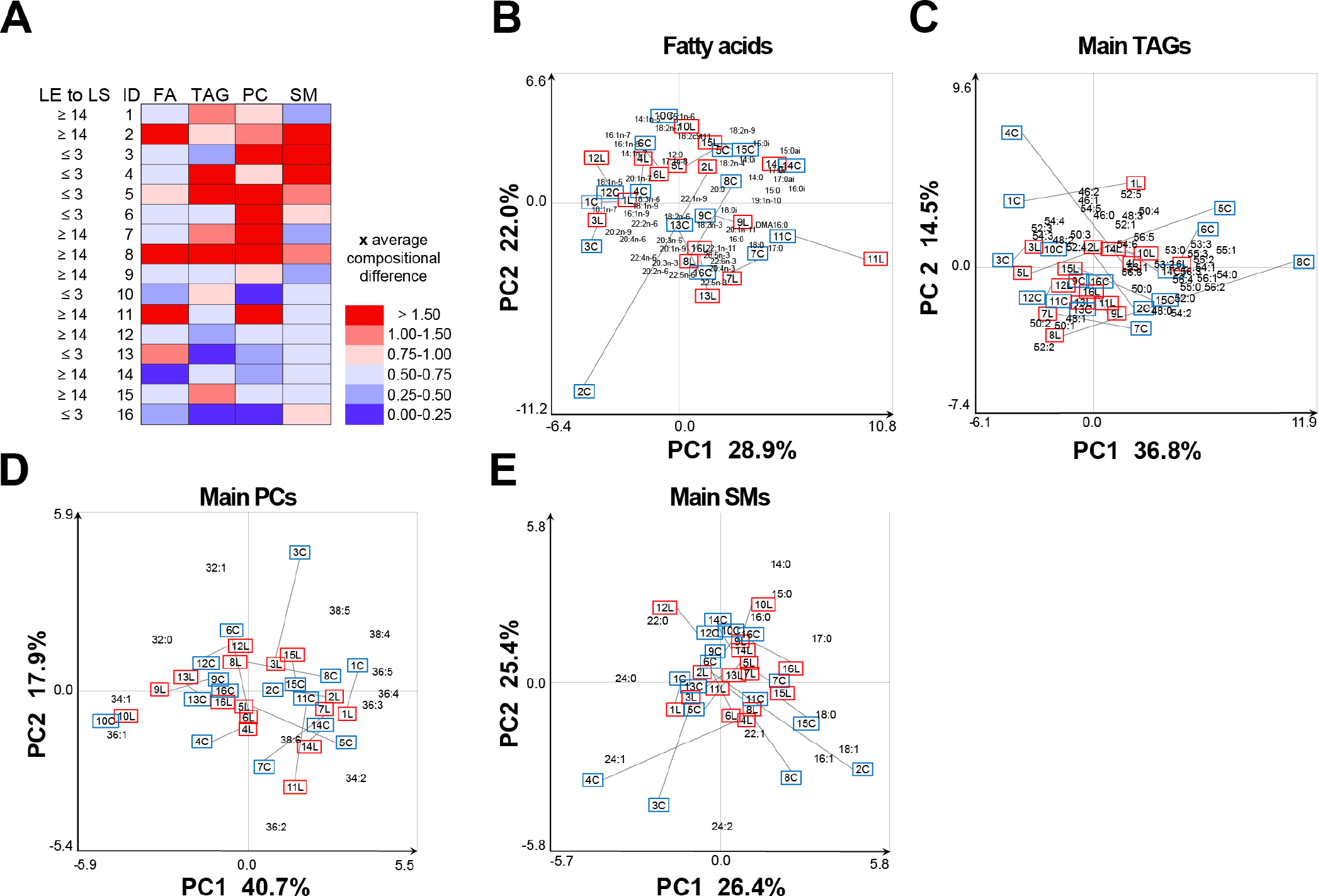
Fatty acid and lipid species profiles of lymphedema arm compared to control arm. (**A**) Duration of lymphedema and compositional difference of the two arms of each individual (numbered 1-16) in the PCA scores using fatty acid (FA), triacylglycerol (TAG), phosphatidylcholine (PC) or sphingomyelin (SM) profile data (mol%) as loadings. The degree of the difference was normalized to the average compositional difference of the two arms in the 16 individuals and illustrated in heatmap format (similar composition = 0, average difference = 1, and the largest compositional differences between the two arms got values >1.5). The map was derived from PCA analysis of the (**B**) FA, (**C**) main TAG species, (**D**) main PC species, and (**E**) main SM species. LE to LS = years from the start of lymphedema to liposuction; ID = ID-numbers of the individuals in the PCA plots B-E. Variable abbreviation: [total number of acyl chain carbons]:[total number of double bonds]. In addition, for FA, the double bond positions were defined by the n-x system, and iso- and anteiso branches were marked by i and ai, respectively. Plasmalogen-derived alkenyl chain derivatized to dimethyl acetal was marked with DMA. On the PCA biplots, the samples were marked with the ID and letter C = control arm (inside blue frame) or L = LE arm (inside red frame).

## DISCUSSION

The multi-omics approach used in this study provides a detailed analysis of the molecular phenotype of LE-induced adipose tissue. The functional genomics analyses demonstrated that inflammatory and angiogenesis pathways are induced in the edematous adipose tissue compared to the healthy arm, whereas pathways related to epidermal differentiation, cell-cell junctions, and water homeostasis were repressed. Interestingly, the gene expression changes in LE-induced adipose tissue expansion were highly different from the ones found in the subcutaneous adipose tissue in patients with obesity. However, the metabolomes and lipidomes did not differ much between the adipose tissues of LE and the healthy arm, but instead, the interindividual variation was much larger. This indicates that the subcutaneous adipose tissue composition is largely determined by the diet and not much affected by LE or its duration.

The whole-genome transcriptomic analysis revealed that the expression of more than 1400 genes underwent changes in the adipose tissue between the affected and non-affected arm. This shows that the LE-induced adipose tissue significantly differs from the same patient’s healthy adipose tissue. We also observed that the affected genes and pathways were highly similar in the short- and long-term LE patients, indicating that the phenotype is induced early during the disease development and does not markedly change thereafter or in response to edema-reducing treatments.

In both LE and obesity, there is a marked expansion of adipose tissue; however, the nature of this growth proved to be highly different. This means that the LE-related adipose tissue is unique, both when compared to healthy adipose tissue and to expanded adipose tissue in obesity; it should therefore be treated differently. In clinics, obesity is a risk factor for developing lymphedema after the surgery, which is caused by the excessive amount of adipose tissue that further impairs lymph transport in the affected arm. We have also found that adipocytes in lymphedematous arms and legs are larger than in the non-affected extremity ^46^. This is in line with the finding that over time larger adipocytes are less insulin sensitive, which we find in the transcriptomics data.

The finding that the metabolomic and lipidomic profiles of the adipose tissue in the lymphedema arm did not differ much from those of the healthy arm was unexpected. Our findings highlight the effect of diet on adipose tissue lipidomic and metabolomic composition even more than the disease, although the transcriptomic landscape was markedly affected by LE. Since the lipid turnover in adipose tissue is on average only 1–4 years ^47^, the pathways of lipid metabolism, adipocyte differentiation, and insulin responses, that were downregulated in the arms after having LE for ≥14 years, had enough time to modify the adipose tissue FA and lipid compositions. Nevertheless, no consistent metabolome or lipidome differences associated with the disease were found between the LE arms and their pairs (except for valine, discussed later). This suggests that the influence of local differences in the metabolic pathways is masked by a similar supply of diet-derived molecules from the systemic pool via circulation.

In most of the studied patients, the lipidome variation within the patients (between diseased and healthy arms) was smaller than the variation between the patients. In addition, the differences between the two arms were qualitatively different in different patients. The only metabolite that was found to be significantly changed was valine, a branched-chain amino acid. Also, other branched-chain amino acids leucine and isoleucine were predicted to be downregulated based on the reporter metabolite analysis, but these amino acids were not detected by the metabolomics platform used in our study. Interestingly, one of the transporters for valine, *SLC6A15* was significantly repressed at mRNA level, indicating that branched-chain amino acid metabolism seems to be changed in the LE adipose tissue. These amino acids have been related to obesity-induced type 2 diabetes ^48^, pointing to a novel direction for further studies in LE. Interestingly, this also goes to the other direction, as increased levels of branched-chain amino acids are found in obesity.

It is likely that the differences we found in the transcriptomics results are caused mainly by other cells than adipocytes. For example, blood and lymphatic endothelial cells, macrophages, and fibroblasts have a large impact on the adipose tissue function and phenotype. An injurious immune response after the surgery and radiation, combined with insufficient reparative immune response, have been proposed to be a major driver for lymphatic vascular dysfunction leading to adipose deposition and fibrosis ^49^. Mechanistic insights have been obtained from the modified mouse tail lymphedema model, where inflammation was shown to precede adipogenesis ^50^. Furthermore, leukotriene B4 antagonism by ketoprofen has been shown to effectively ameliorate experimental lymphedema in mice ^51^.

A recent study suggested that adipose tissue in LE patients has increased basal lipolysis and cytokine production compared to adipose tissue from healthy control subjects ^52^. A major strength of the present study is the comparison of LE adipose tissue to healthy adipose tissue of the same patient. We further identified patients with short- and long-term disease, allowing us to compare the adipose tissue responses in earlier versus prolonged phases of the disease. We also had the opportunity to compare the findings to a large dataset from female subjects with obesity. As a limitation, we were able to run the omics studies in a limited number of patients. For validation of the possibility to use for example *CETP*, which was the most upregulated gene, as a biomarker for LE, it would have been useful to have serum samples from these patients. Thus, further studies are needed to address if CETP protein could serve as a serum marker for those breast cancer patients, who are likely to develop LE and related adipose tissue accumulation.

This study provides the first comprehensive map of transcriptional changes that change during LE development over time. An increased understanding of the molecular characteristics of the LE-induced adipose tissue expansion may provide new avenues for the development of novel therapies for LE patients.

## Authors’ contributions

S.K., K.A., R. Kivelä conceptualized the study; S.K. and R. Kivelä performed investigation and wrote the original draft of the manuscript. H.B. provided the patient samples. S.K., S.L., C.Z., R. Käkelä, A.M., R. Kivelä contributed to data curation, data analysis; S.K., S.L., C.Z., R. Käkelä, R. Kivelä contributed to methodology and data visualization; S.L., C.Z., R. Käkelä, A.M., H.B., and K.A. reviewed and edited the manuscript. M.R.T., H.B., and K.A. provided resources. K.A. provided supervision and acquired funding. All authors read and approved the final manuscript.

## DATA AVAILABILITY

Data are available upon reasonable request. The differential gene expression comparisons of LE versus control arms in short- and long-term LE can be found in **supplemental data table 1**.

## ACKNOWLEDGEMENTS

We gratefully acknowledge Dr. Bushra Shaida for the preparation of adipose tissue samples (Lund University); Mari Jokinen for RNA isolation, qPCR, and metabolomics sample preparations (Wihuri Research Institute); and Madeleine H. Lackman for help with circos plots (University of Helsinki). Bulk RNA isolation and sequencing was performed at the BEA Core Facility (Karolinska Institute). Metabolomics analyses were carried out at the Biocenter Finland and HiLIFE supported Metabolomics Unit (Institute for Molecular Medicine Finland FIMM, HiLIFE, University of Helsinki). This work was funded by the Leducq Foundation Transatlantic Network of Excellence Lymph Vessels in Obesity and Cardiovascular Disease (grant 11CVD03 to K.A.), Jenni and Antti Wihuri Foundation, and the Novo Nordisk Foundation (NNF16OC0023554). S.K. was supported by the Swiss National Science Foundation (Advanced Postdoc.Mobility grant number: P300PB_164732), Orion Research Foundation, Finnish Foundation for Cardiovascular Research, Maud Kuistila Memorial Foundation, Academy of Finland (330053) and University of Helsinki. R. Kivelä. was supported by the Academy of Finland (297245), the Sigrid Jusélius Foundation and the Finnish Foundation for Cardiovascular Research. H.B. was funded by the Swedish Cancer Society, Stockholm (CAN 2016/639, CAN 2013/505) and Skåne County Council’s Research, and Development Foundation (REGSKANE-454091, REGSKANE-626131, REGSKANE-816751, Project 2020-0460).

